# Genome-wide identification, characterization and expression analysis of the Bcl-2 associated athanogene (BAG) gene family in *Physcomitrium patens*

**DOI:** 10.1101/2020.12.23.424083

**Authors:** Alexanda Castro, Laura Saavedra, Cecilia Ruibal, Ramiro Lascano, Sabina Vidal

## Abstract

The Bcl-2-associated athanogene (BAG) family is an evolutionarily conserved, multifunctional group of co-chaperones regulators that modulate a number of diverse processes. Plant *BAG* genes were identified to play an extensive role in processes of programmed cell death (PCD) ranging from growth and development to stress responses. In this study, we identified *BAG* genes from different photosynthetic organisms in order to gather evolutionary insights on these proteins followed by an *in silico* characterization of the *BAG* family in the bryophyte *Physcomitrium patens*. Ten putative *PpBAGs* harbouring a characteristic BAG domain were grouped into two subfamilies based on the presence of additional conserved domains and phylogenetic distances. Group I consisted of PpBAG4 and PpBAG5, containing an additional ubiquitin-like domain, and PpBAG10 with only the BAG domain. Group II consisted of PpBAG1–3 and PpBAG6-9, containing a calmodulin-binding IQ motif, a novel feature associated with plant BAG proteins. Interestingly, PpBAG9 exhibits an EF-Hand domain, not reported to date in this class of proteins. Caspase cleavage sites in PpBAG1, PpBAG3, PpBAG4-5 and PpBAG9 were predicted. *In silico* analysis of *BAG* genes revealed the presence of stress responsive elements, and a stress-regulated expression pattern which appears to be dependent on specifically organized promoter regulatory elements. According to our analyses, the present data suggest that some members of *P. patens BAG* gene family may play a role in heat responses, autophagy and pathogen immunity. Further studies are required to unveil the role of specific members of this gene family in PCD and stress responses in *P. patens*.

**Key message:** Genome-wide identification and phylogenetic relationships combined with *in silico* gene-expression profiling and protein-interaction analysis of PpBAGs in *Physcomitrium patens*, highlight the importance of a particular set in stress tolerance.

## Introduction

Programmed cell death (PCD) is essential to maintain tissue homeostasis in many life forms. In plants, in the same way as in animals, PCD is the mechanism by which a series of physiological processes such as seed development, germination, growth and differentiation, reproduction and senescence are regulated. PCD also plays an important role in other processes such as resistance to pathogens and to unfavorable environmental conditions. The molecular mechanisms of plant PCD are less understood than the mechanisms controlling this process in animal systems. Although there is a certain degree of parallelism between the pathways operating in plant and animal PCD, the existence of apoptotic type of PCD in plants is still a matter of debate. However, in recent years, important evidence has emerged to support the presence of such a process in plants (Reape *et al*., 2008; Daneva *et al*., 2016; Balakireva *et al*., 2019).

Structural analyses have enabled the identification of the B cell lymphoma 2 (Bcl-2)-associated athanogene (BAG) family of co-chaperones in plants, one of the core conserved regulators of PCD in animal apoptosis (Thanthrige *et al*., 2020). *BAG* genes are an evolutionarily conserved family with homologs found from yeast to animals, including plants (Froesch *et al*., 1998; Emanuelsson *et al*., 2000; Sondermann *et al*., 2001; Kim *et al*., 2002, Kabbage and Dickman, 2008). The first *BAG* gene discovered was the human *BAG-1*, which encodes a protein that can interact with Bcl-2 to greatly enhance the antiapoptotic effect of Bcl-2. BAG proteins are characterized by a conserved region located near the C terminus, termed the BAG domain (BD). The BD domain contains 110–130 amino acids and is comprised of three α helices of 30–40 amino acids each; where the second and third helices mediate direct interaction with the ATPase domain of Hsp70/Hsc70 molecular chaperones (Sondermann *et al*., 2001). The human BAG family has six members which are involved in the modulation of numerous physiological processes, such as apoptosis, tumorigenesis, neuronal differentiation, stress responses and cell-cycle progression (Kabbage and Dickman, 2008; Behl 2016). These proteins are known to regulate positively or negatively the function of Hsp70/Hsc70, and to form complexes with a different range of transcription factors (Terada *et al*., 2000).

Plant genomes also encode BAG proteins (Doukhanina *et al*., 2006; Kang *et al*., 2006; Thanthrige *et al*., 2020); however, only a few plant BAG proteins have been studied, revealing a role in cytoprotection under stress conditions (Thanthrige et al., 2020). *Arabidopsis thaliana* and *Oryza sativa* genomes contain seven and six homologs of the *BAG* gene family, respectively (Doukhanina *et al*., 2006; Rana *et al*., 2012). The domain organization of four AtBAGs (AtBAG1-4) was found to be similar to their animal homologs, while the remaining three proteins (AtBAG5-7) contain a calmodulin-binding domain, the isoleucine-glutamine (IQ) motif near the BAG BD. The presence of an IQ motif is a novel feature associated with plant BAG proteins reflecting possible divergent mechanisms involved in plant-specific PCD (Doukhanina et al., 2006). Although several *BAG* family members have been characterized in tracheophytes (vascular plants), the function of these genes has not been characterized in non-vascular species such as the moss *Physcomitrium patens (P. patens*, previously known as *Physcomitrella patens*). Bryophytes, comprising mosses, liverworts and hornworts originated around 500 million years ago, and as such, they can provide insight into the adaptive mechanisms that allowed the transition from water to land (Kenrick, 2017; Rensing, 2018). Among bryophytes, mosses are the most diverse group and have been studied since the 1930s. During the past two decades, the moss *P. patens* has emerged as a very useful model system for the analysis of many aspects of plant biology and evolution. *P. patens* genome has been sequenced and annotated based on sequence homology to known genes and domains and offers a valuable resource for research aimed at understanding functional evolution in plants (Rensing *et al*., 2008).

In this work, we present the first analysis of the *BAG* gene family from a bryophyte, as a way to better understand BAGs evolution in land plants. To gain further insight into the possible functions of moss *BAG* genes, we carried out a comprehensive analysis of the *BAG* gene family. We studied the structural properties, gene structure and organization, phylogeny, and expression profiles of the *BAG* gene family in different tissues and under various hormonal and stress treatments. Our analysis of *P. patens BAG* genes at the genomic level provides a solid basis for further functional characterization of specific genes from this family.

## Material and Methods

### Identification and phylogenetic analysis of BAG proteins

Sequences containing the BAG domain from 14 algae and plant species (Suppl. Table S1) namely, *Micromonas sp. RCC299, Micromonas pusilla CCMP1545, Coccomyxa subellipsoidea C-169, Chlamydomonas reinhardtii, Volvox carterii, Physcomitrium patens, Sphagnum fallax, Marchantia polymorpha, Selaginella moellendorffii, Arabidopsis thaliana, Glycine max*, and *Oryza sativa* were retrieved combining different methods: from the Phytozome v12.1 database (https://phytozome.jgi.doe.gov) through the BioMart data query tool (https://phytozome.jgi.doe.gov/biomart/martview) using the BAG domain BD PFAM accession term PF02179 (http://pfam.sanger.ac.uk/), through a BLASTP search using the BAG-domain amino acid sequence from different BAG proteins as query, and also using the profile hidden Markov Model tool (HMMER) with the BAG protein domain in Ensembl protein sequence databases.(https://www.ebi.ac.uk/Tools/hmmer/search/phmmer). BAG genes from *Klebsormidium nitens* v1.1 were retrieved from the *Klebsormidium nitens* project (http://www.plantmorphogenesis.bio.titech.ac.jp/~algae_genome_project/klebsormidium/) through the BLASTP search using the BAG amino acid sequences from *A. thaliana* (Doukhanina *et al*., 2006). The presence of different domains was further confirmed at the NCBI CCD search domain (https://www.ncbi.nlm.nih.gov/Structure/cdd/wrpsb.cgi), PFAM (https://pfam.xfam.org) and SMART (https://smart.embl-heidelberg.de) databases.

Translated protein sequences were aligned with MUSCLE (Mega 7.0 version, Tamura et al., 2011). The evolutionary history was inferred using the Neighbor-Joining method and the bootstrap consensus tree was inferred from 1000 replicates. The evolutionary distances were computed using the Poisson correction method and are presented in the units of the number of amino acid substitutions per site.

Genomic schematic diagrams of the BAGs were obtained by comparing the genomic sequences and their predicted coding sequences using the GSDS tool (http://gsds.gao-lab.org/) (Hu *et al*., 2015).

### Physicochemical properties and protein structure model of *PpBAG* genes

PpBAG protein sequences were analyzed using the ExPASy-ProtParam tool: http://web.expasy.org/protparam/ to calculate the number of amino acids, molecular weight, and theoretical pI. (Gasteiger *et al*., 2005). ExPASy Peptide Cutter tool (Wilkins *et al*., 1999) was used to search potential protease cleavage sites in PpBAGs proteins. The predicted subcellular localization of the PpBAG protein family was analyzed using GTP-Pp, a prediction tool suitable for sequences from *P. patens* (Fuss *et al*., 2013).

The homology models for *P. patens* BAG proteins were generated through SWISS-MODEL web server. Each model was generated based on automated identification of template model. Visualization was done in Pymol 2.4.0 (DeLano, 2002)

### Putative cis-Regulatory Elements in *BAG* Genes

To identify putative cis-regulatory elements in the promoter region of *PpBAG* genes, a 1.5 kb fragment upstream of the translation initiation codon of each *PpBAG* was extracted using Phytozome. Sequences were then analyzed using Plant cis-acting regulatory DNA elements (PLACE; http://www.dna.affrc.go.jp/PLACE/), Plant cis-acting regulatory elements (PlantCARE; http://bioinformatics.psb.ugent.be/webtools/plantcare/html/, and PlantPAN (http://plantpan.itps.ncku.edu.tw/). Results were filtered to keep the number and detailed position of cis-element regarding growth and development, abiotic stresses and hormones responses in plant.

### *In silico* expression analysis of *BAG* genes

The expression profiles of *P. patens* BAG genes during plant development and stress conditions were analyzed in silico using microarray experiments data from Genevestigator database (https://genevestigator.com/) (Hruz et al., 2008), *Physcomitrella* eFP database (http://bar.utoronto.ca/efp_physcomitrella/cgi-bin/efpWeb.cgi) (Ortiz-Ramírez *et al*., 2016) and PTEATmoss (https://peatmoss.online.uni-marburg.de/index) (Fernandez-Pozo *et al*., 2020).

### Protein-protein interaction analysis

The Protein-protein interaction (PPI) networks were identified in *P. patens* using the STRING search tool (version 11.0) (http://string-db.org) with default parameters. Sources in STRING include experimentally determined interactions, curated databases, and information of co-expression, fusion, text mining and co-occurrence (Szklarczyk *et al*., 2019).

## Results

### Identification, phylogenetic relationships and gene structure analysis of *BAG* genes in Viridiplantae

In order to identify BAG proteins from different photosynthetic organisms, the sequence of the conserved BAG domain was used as a query to search in a number of algae and plant genomes form Ensembl protein sequence databases using the profile hidden Markov Model tool (HMMER). In addition, BLASTN and BLASTP searches were performed using the BD amino acid sequence from different BAG proteins as a query in Phytozome v12.1 database, or through the BioMart data query tool using the BAG domain PFAM accession term PF02179. The search was first performed in *Arabidopsis thaliana* resulting in seven AtBAGs, previously described by Doukhanina *et al*., (2006). Subsequently, the same strategy was used to search within genomes of chlorophytes (*Micromonas sp. RCC299, Micromonas pusilla CCMP1545, Coccomyxa subellipsoidea C-169, Dunaliella salina*), bryophytes (*Physcomitrium patens, Sphagnum fallax, Marchantia polymorpha*), lycophytes (*Selaginella moellendorffii*) and other angiosperms (*Glycine max* and *Oryza sativa*). BAG protein sequences from the charophyte *Klebsormidium nitens* v1.1 were retrieved from the *Klebsormidium nitens* project (see Materials and Methods). As a result, 74 BAG domain-containing proteins were identified and used in this study (Suppl. Table S1). The presence of the BAG domain and additional domains exhibited by these proteins were further analyzed by the NCBI CCD search domain, PFAM and SMART databases.

The phylogenetic relationship between BAG proteins from the different photosynthetic organisms ranging from streptophyte algae to angiosperms resulted into 2 groups: Group I containing members with a ubiquitin-like (UBL) domain in addition to the BD domain; and Group II including proteins with a Calmodulin (CaM)-binding module (IQ motif), located towards the N-terminal side of the BD domain (Fig. 1a). This classification was previously described for the *A. thaliana* and *O. sativa* BAG protein families (Doukhanina et al., 2006; Rana et al; 2012) and this notion is now extended through the analysis of the BAGs distribution across different species in Viridiplantae. Interestingly, our analysis revealed two subfamilies within Group I: IA and IB. The first includes the most ancestral BAGs, containing BAGs with an UBL domain from the chlorophytes *Micromonas* sp. RCC299, *Micromonas pusilla* CCMP1545, *Coccomyxa subellipsoidea* C-169, and the charophyte *K. nitens* (kfl00722_0030), and also include members which harbor a BAG domain alone from *K. nitens* (kfl00722_0040) and the bryophytes *P. patens* (PpBAG10), *M. polymorpha* (Mapoly0049s0106) and *S. phallax* (Sphfalx0219s0026). The genomes from all chlorophytes analyzed contain single BAG genes, whereas the charophyte *K. nitens*, contains two BAG members: one harboring a UBL domain plus the BD domain, and another with a single BD. All members of Group IB contain BAG proteins with the UBL domain in addition to the BD domain. On the other hand, group II harbors members with a CaM-binding IQ motif, which is present exclusively in land plant BAGs suggesting plant specific modes of signaling and/or regulation of BAGs by Ca^+2^. None CaM-binding motifs were found in the BAG proteins from the streptophyte algae analyzed, suggesting that the acquisition of this domain in BAG proteins has evolved with the terrestrialization of plants, whereas BAG proteins containing an UBL domain evolved earlier. One member of this group, PpBAG9, has an EF-Hand motif near the C-terminal region, a novel feature that has not been described so far for members of BAG proteins (Fig. 2).

**Fig. 1.**
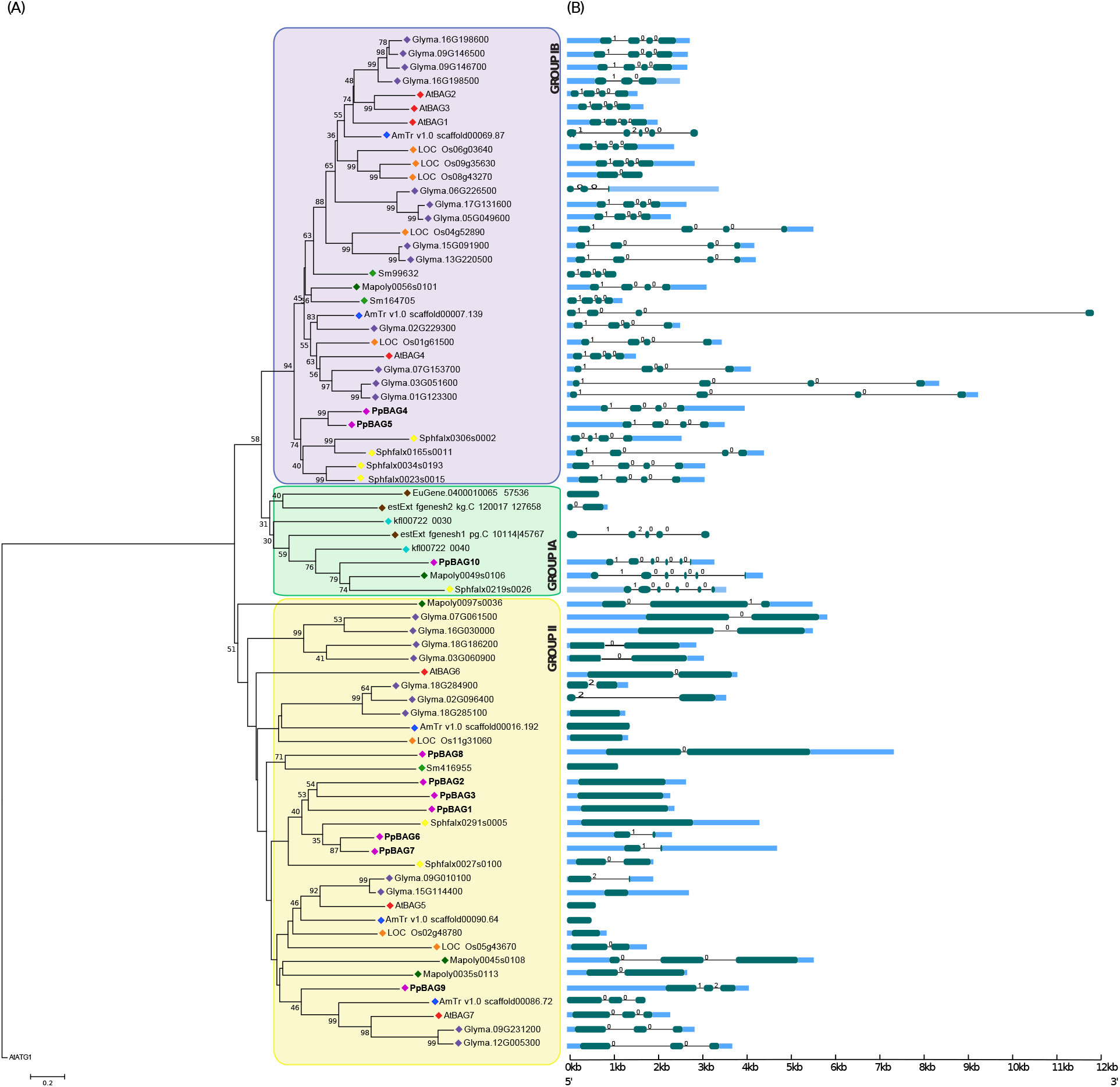
Phylogenetic tree and exon-intron analysis of the BAG family and gene structure in selected Viridiplantae members. **a** The evolutionary history of BAG proteins was inferred using the Neighbor-Joining method. The percentage of replicate trees in which the associated taxa clustered together in the bootstrap test (1000 replicates) are shown next to the branches. The tree was rooted using the AtATG1 protein as an outgroup. The coloured rhombuses indicate the following or species: brown, chlorophytes (*Micromonas sp. RCC299, Micromonas pusilla CCMP1545, Coccomyxa subellipsoidea C-169*); turquoise, *Klebsormidium nitens;* purple, *Physcomitrium patens;* yellow, *Sphagnum fallax;* dark green, *Marchantia polymorpha;* light green, *Selaginella moellendorffii;* blue, *Amborella trichopoda;* red, *Arabidopsis thaliana*; orange, *Oryza sativa;* violet, *Glycine max*. **b** Exon-intron structures were described using GSDS 2.0. Green, round-cornered rectangles represent exons, black lines represent introns, and blue sky rectangles represent untranslated regions (UTRs). The numbers 0, 1, and 2 represent intron phases.

**Fig. 2.**
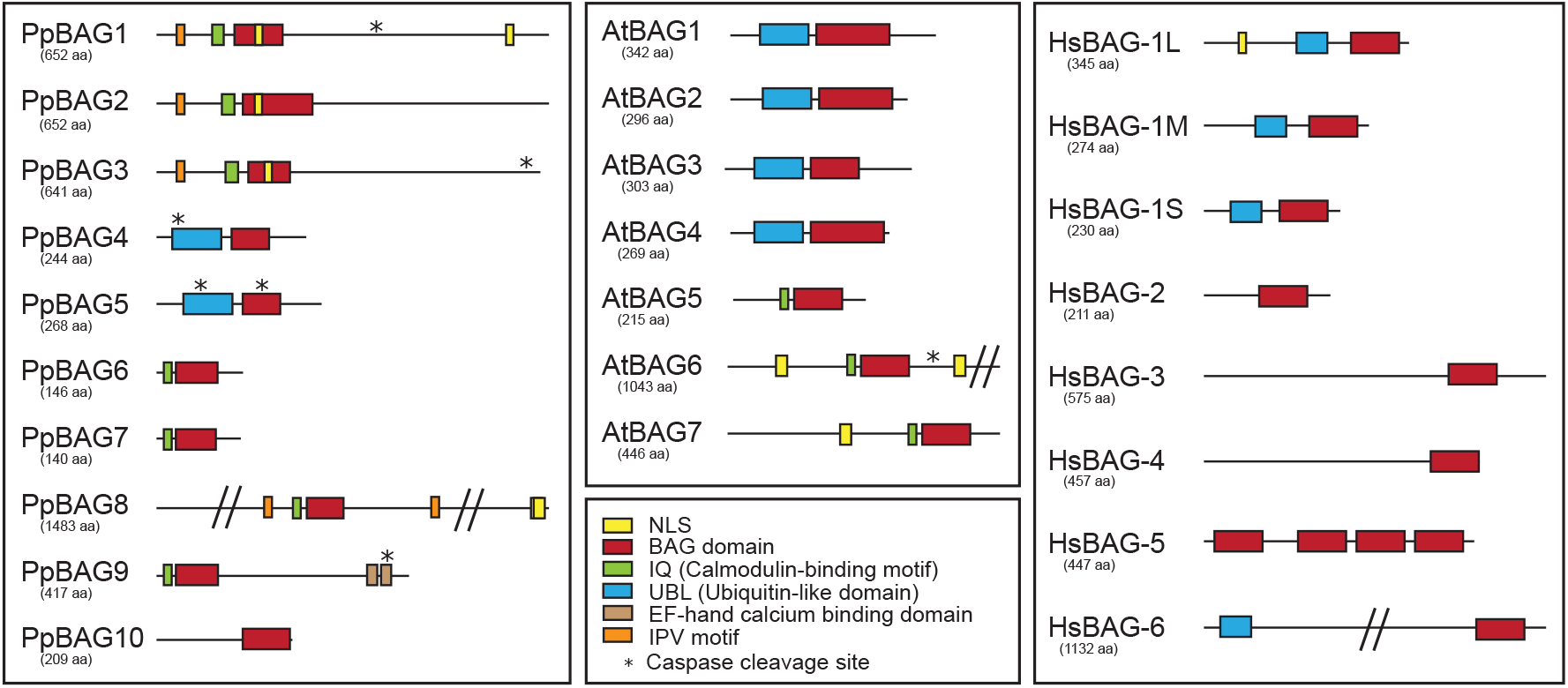
Domain sequences of PpBAGs from *P patens*, identified by SMART database (http://smart.embl-heidelberg.de), compared with human (Hs) and *Arabidopsis thaliana* (At) BAG proteins. The red, green, blue, orange, brown and yellow squares indicate the BAG domain, the calmodulin-binding motif (IQ), the ubiquitin-like domain (UBL), the IPV motif, the EF-hand calcium binding domain and the nuclear localization signal, respectively. Putative caspase cleavage sites are shown with an asterisk. Protein length in amino acids is displayed under each BAG.

In the case of *P. patens*, our analysis revealed the existence of ten genes encoding BAG proteins (PpBAGs) which are listed in Table 1. *PpBAG* genes are dispersed throughout the *P. patens* genome, being *PpBAG2, PpBAG4* and *PpBAG10* located on chromosome 26, *PpBAG3* and *PpBAG5* on chromosome 5, *PpBAG7* and *PpBAG8* on chromosome 19, and *PpBAG1, PpBAG6* and *PpBAG9* on chromosomes 13, 21 and 7, respectively. The different PpBAG proteins were grouped into the two groups as follows: PpBAG10 in Group IA, PpBAG4 and PpBAG5 in Group IB, and PpBAG1-3 and PpBAG6-9 in Group II (Figs. 1 and 2).

**Table 1.**
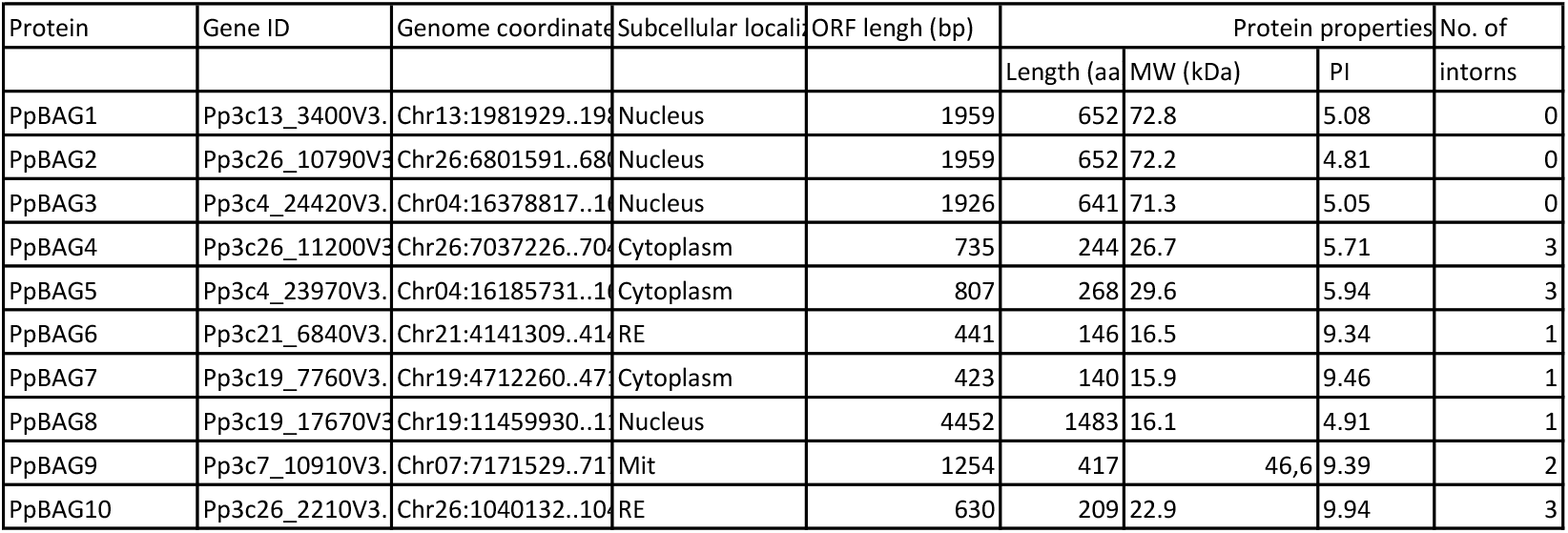
Properties of predicted BAG proteins in *P. patens*.

| Protein | Gene ID | Genome coordinate | Subcellular localiz | ORF length (bp) | Protein properties |  |  | No. of |
| --- | --- | --- | --- | --- | --- | --- | --- | --- |
|  |  |  |  |  | Length (aa) | MW (kDa) | PI | intorns |
| PpBAG1 | Pp3c13_3400V3. | Chr13:1981929..198 | Nucleus | 1959 | 652 | 72.8 | 5.08 | 0 |
| PpBAG2 | Pp3c26_10790V3. | Chr26:6801591..680 | Nucleus | 1959 | 652 | 72.2 | 4.81 | 0 |
| PpBAG3 | Pp3c4_24420V3. | Chr04:16378817..16 | Nucleus | 1926 | 641 | 71.3 | 5.05 | 0 |
| PpBAG4 | Pp3c26_11200V3. | Chr26:7037226..703 | Cytoplasm | 735 | 244 | 26.7 | 5.71 | 3 |
| PpBAG5 | Pp3c4_23970V3. | Chr04:16185731..16 | Cytoplasm | 807 | 268 | 29.6 | 5.94 | 3 |
| PpBAG6 | Pp3c21_6840V3. | Chr21:4141309..414 | RE | 441 | 146 | 16.5 | 9.34 | 1 |
| PpBAG7 | Pp3c19_7760V3. | Chr19:4712260..471 | Cytoplasm | 423 | 140 | 15.9 | 9.46 | 1 |
| PpBAG8 | Pp3c19_17670V3. | Chr19:11459930..11 | Nucleus | 4452 | 1483 | 16.1 | 4.91 | 1 |
| PpBAG9 | Pp3c7_10910V3. | Chr07:7171529..717 | Mit | 1254 | 417 | 46,6 | 9.39 | 2 |
| PpBAG10 | Pp3c26_2210V3. | Chr26:1040132..104 | RE | 630 | 209 | 22.9 | 9.94 | 3 |

The structural diversity generated by losses or gains of introns within gene families is one of many evolutionary mechanisms that create genetic diversity (Doolittle, 1978; Gilbert, 1987; Gilbert, 1997). In order to understand the possible structural relationship among *BAG* gene orthologues and paralogues, the exon/intron organization of the different *BAG* genes was analyzed using the GSDS 2.0 software. *PpBAG4* to *PpBAG10* contain introns, whereas *PpBAG1-3* are intronless genes (Fig. 1b). Intron positions and intron phases of the *BAG* genes were found to be well-conserved in Group IB. (Fig. 1b). However, Group IA and Group II exhibited more diversity. It has been speculated that lineage-specific intron loss events might have occurred during the expansion and structural evolution of genes, thereby generating gene structural diversity (Nguyen *et al*., 2006). The intron phases analysis showed that the 3 introns of the gene exon/intron structure of the 33 genes analyzed from the Group IB were all conserved and share the same pattern of phases, with the exception of two genes from soybean (*Glyma.16G198500, Glyma.06G226500*), one from rice (*Os08g43270*) and one from *Amborella (AmTr v1.0 scaffold00069.87*). The first intron showing a phase-1 is located in the portion corresponding to the N-terminal domain of the protein, the most diverse region. The following two introns, in phase-0, were located in more conserved regions, e.g., in the middle and in the C-terminal domain of the protein (Fig. 1b). These results are consistent with those previously reported by (Rogozin *et al*., 2012), showing that introns with phase-0 are usually located in more conserved portions of genes than introns with phases 1 or 2.

### Motif analyses and predicted caspase cleavage sites of BAG proteins in *P. patens*

The ten deduced *P. patens* BAG proteins have varying theoretical isoelectric points, ranging from 4.81 (PpBAG2) to 9.46 (PpBAG7), as well as predicted molecular weights, from 15.9 kDa (PpBAG7) to 161.7 kDa (PpBAG8). The *in silico* subcellular locations depicted for these proteins were nuclear for PpBAG1-3 and PpBAG8; cytoplasmic for PpBAG4, PpBAG5 and PpBAG7; endoplasmic reticulum for PpBAG6 and PpBAG10, and mitochondrial for PpBAG9 (Table 1).

*P. patens* BAGs exhibit a conserved modular structure compared to their orthologs from other plant species. The UBL domain present in PpBAG4 and PpBAG5 maintains the conserved lysine of the ubiquitin_like family domains, one of seven lysines involved in chain linkage in ubiquitin. This feature is shared with all the BAG proteins containing a UBL domain analyzed in this study, with the exception of *M. pusilla* BAG (Fig. 3b). In addition, alignment of the IQ motif present in *P. patens* and *Arabidopsis* BAGs showed a general aa conservation within the IQ consensus sequence IQXXXRGXXXR (Putkey *et al*., 2003) (Fig. 3a). Previous analysis of AtBAG5 showed that the IQ-motif preferentially binds the calcium-free state of Calmodulin, apocalmodulin, (apo-CaM) and the BAG domain binds with Hsc70. In the presence of Ca^2+^, Ca^2+^-bound CaM alters its binding mode to AtBAG5 by disrupting AtBAG5/Hsc70 binding, inducing the release of Hsc70 which in turn suppress ROS generation, and consequently delaying leaf senescence (Luhua Li *et al*., 2016).

**Fig. 3.**
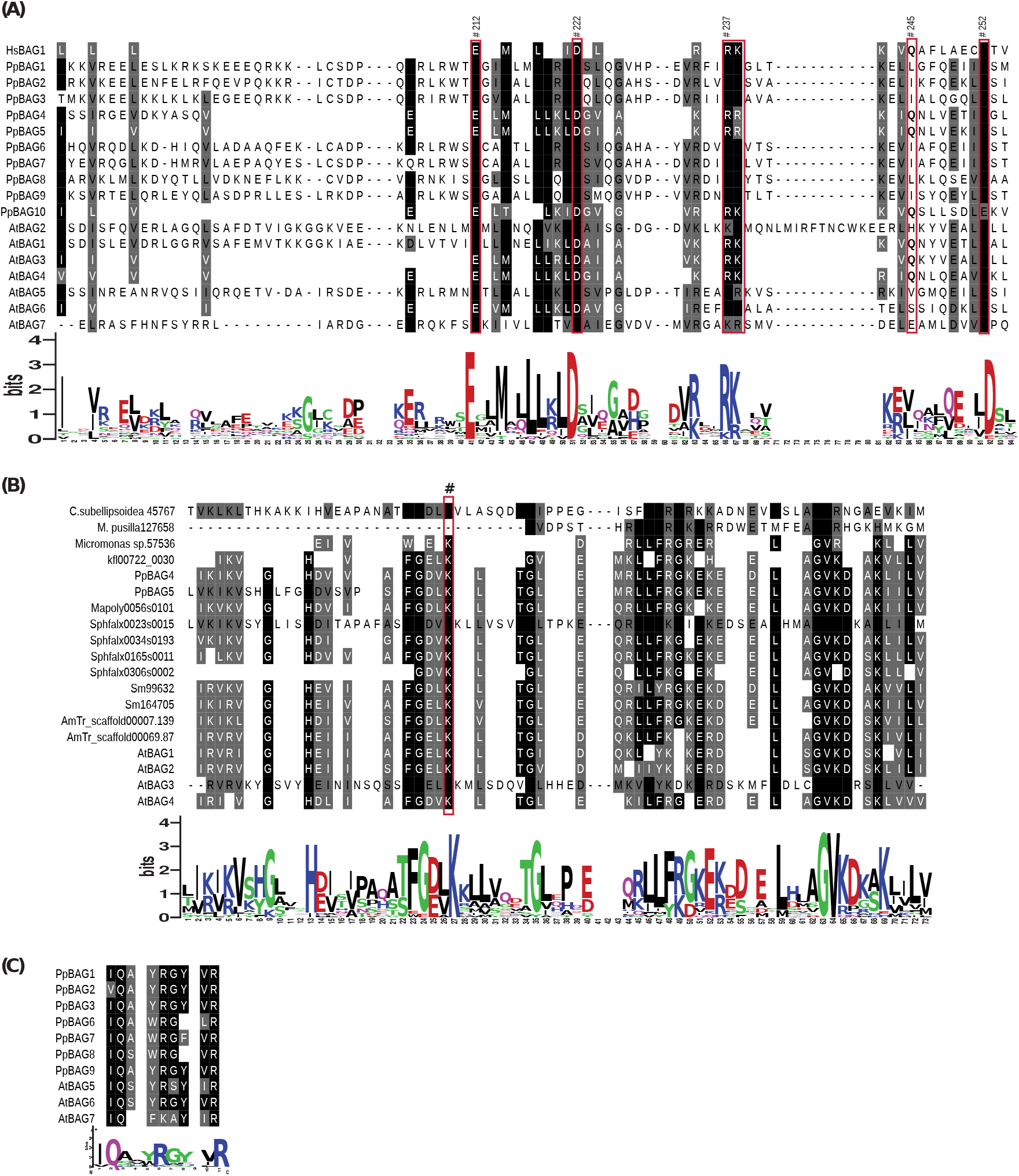
Alignment of amino acid sequences of the different domains of BAG proteins and their corresponding sequence logos determined by the WebLogo program. **a** Alignment of the BAG domain from HsBAG1, PpBAGs and AtBAGs. The numbers correspond to HsBAG1 conserved residues forming the interaction surface with the Hsc70 ATPase domain. Conserved amino acid residues are shown in black background (identical) or grey background (with > 70% identity). **b** Alignment of the Ubiquitin domain. **c** Alignment of the IQ motif of *P. patens* BAGs.

In the *P. patens* BAG protein family, three members (PpBAG4-5 and PpBAG10) have a domain organization similar to that of mammalian BAG proteins. The resolved crystal structure of the HsBag-1M domain in complex with the ATPase domain of Hsc70 demonstrated that the residues Glu^212^, Asp^222^, Arg^237^ and Gln^245^ are essential to mediate electrostatic interactions for the binding of the monomeric BD domain to the ATPase domain (Sondermann *et al*., 2001). Comparative sequence alignment between the BD domain of the human BAG-1 and the *P. patens* and *Arabidopsis* BAG proteins showed that most of the residues required for Hsp70 binding in mammals are conserved with the exception of Gln^245^ in PpBAG1-3 and PpBAG6-9 and AtBAG2, AtBAG5-7 (Fig. 3a). The three-dimensional structures of the BD domain for the ten PpBAG proteins were predicted through the SWISS-model server, using the *Homo sapiens* BAG4 BD (PDB ID 1M7K) 3-D structure as a template, based on its percentage of highest identity (~22 to 30%) with the PpBAG BDs. The surface of the reported BD solution NMR structure of HsBAG4 with a three-dimensional model of the BD of PpBAG1 to PpBAG10 is shown in Fig. 4. The HsBAG-BD protein exhibits high similarities with respect to charged residue distributions surface residues of the PpBAG4 and PpBAG6-7, while less similarity to PpBAG1-3, PpBAG5 and PpBAG8-10.

**Fig. 4.**
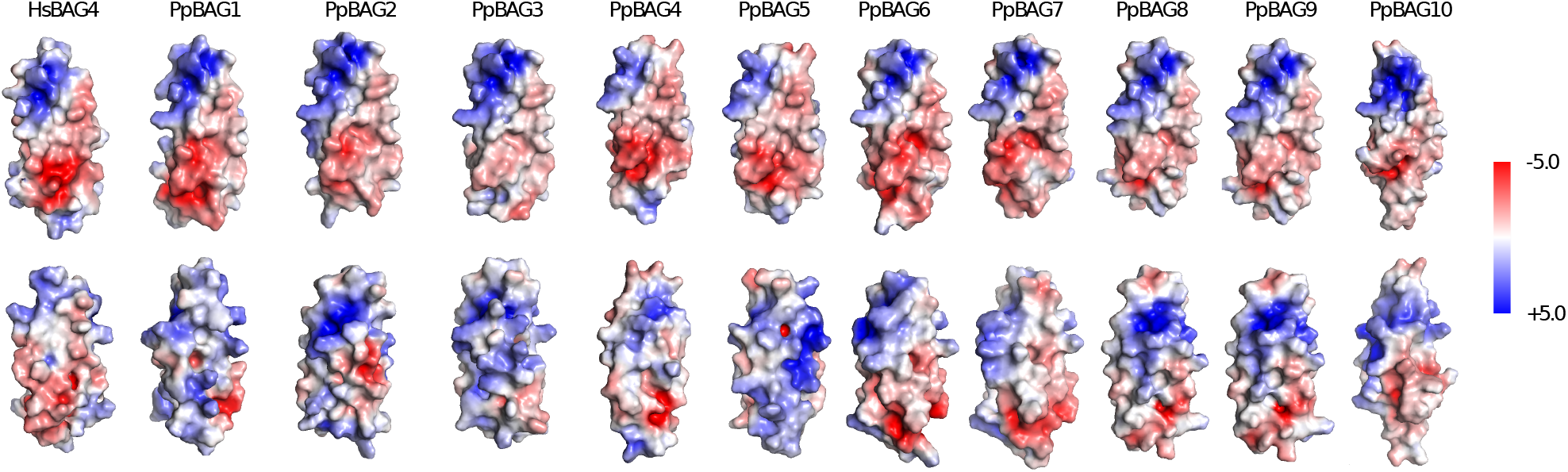
Homology models of the ten *P. patens* BAG BDs as compared to the human BAG4 BD (PDB ID: 1M7K). Positively charged regions are shown in blue, negatively charged regions in red, the upper row represents the front and the bottom row represents a back view of the 3D model for each protein. The structures were predicted by using SWISS-MODEL web server (http://swissmodel.expasy.org/) and processed by PyMol viewer program.

It has been previously reported that resistance to apoptosis is correlated with a reduced caspase activity and enhanced expression of anti-apoptotic proteins like BAGs (Ding *et al*., 2000). To investigate whether PpBAGs contain potential caspase or other protease cleavage sites, the deduced proteins were analyzed using the PeptidCutter program provided by the ExPasy Server. Predicted caspase target sites were identified in PpBAG1, PpBAG3-5 and PpBAG9, suggesting that at least these members of the *P. patens* BAG protein family may undergo processing for cell death/cytoprotective activity (Fig. 2).

### Cis-acting regulatory elements in promoter regions of *PpBAG* genes

A large number of conserved cis-elements are known to regulate gene expression in plant Promoter sequences up to 1.5 kbp upstream from the translation initiation site of each *BAG* gene of *P. patens* were analyzed using different databases for the identification of cis-acting regulatory elements (CAREs). Fig. 5 shows the presence of multiple stress-related and hormonal CAREs in the promoter regions of *P. patens BAG* genes. Elements associated with defense responses (STRE and TC-rich repeats), drought (DRE, MBS), heat shock (HSE), low temperature (LTR), wounding (W-box, WRE3, WUN-motif) and anaerobiosis (AREs) were observed. Among those, the related to drought and defense were the most common together with the binding set of the MYB transcription factor (MYB sequence). Several motifs responding to phytohormones were revealed, including those regulating responses to auxin (TGA-element), salicylic acid (TCA-element, TGA-Box), abscisic acid (ABRE), ethylene (ERE) and methyl jasmonate (CGTCA-motif, TGACG-motif), being those elements that respond to MeJA and ABA the most frequent (Fig. 5). In addition, the TFs binding sites identified in the promoter region of *PpBAG* genes were those associated mainly with stress responses but also with plant growth and developmental processes, such as Myb, bZIP, HSF, WRKY and AP2, C2H2, AT-Hook, bHLH, SBP, Dof, NAC and Homeobox (HD-ZIP), respectively (Table S2).

**Fig. 5.**
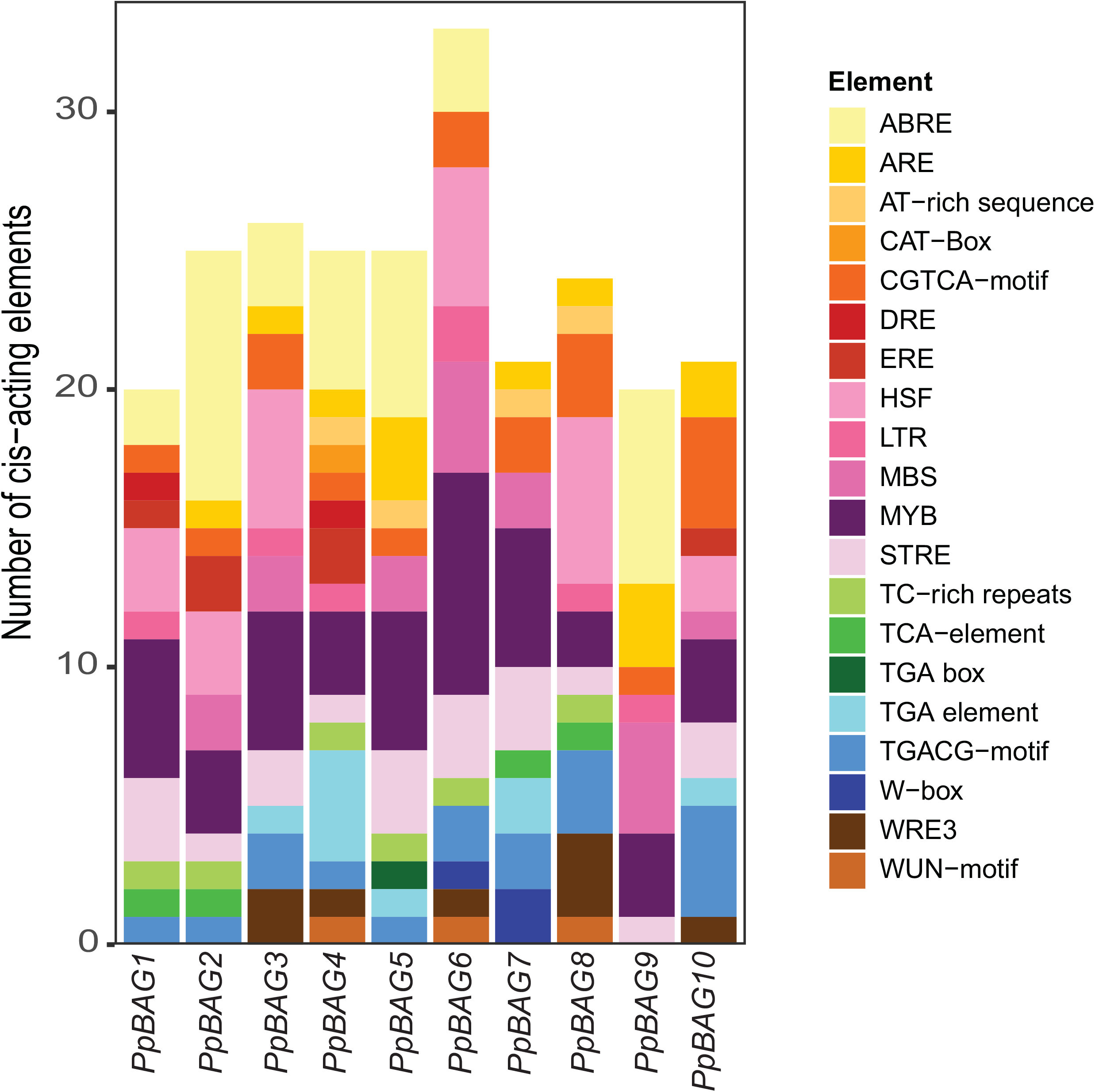
Distribution of stress-related and phytohormone-responsive *cis*-acting elements in the promoters of the *PpBAG* gene family. The cis-acting elements were identified by PlantCARE using the upstream 1500 bp sequences of the *PpBAG* genes. Stress response: ARE, AT-rich sequence, DRE, HSF, LTR, MBS, MYB, STRE, TC-rich repeats, W-box, WRE3 and WUN-motif; Hormone response: abscisic acid (ABRE), ethylene (ERE), auxin (TGA box, TGA element), salicylic acid (TCA-element), methyl jasmonate (CGTCA-motif, TGACG-motif), Growth & Development (CAT Box).

The results suggest that *P. patens BAG* family genes are involved in a variety of stress and plant hormonal responses, and therefore are likely to play a role in essential biological processes such as stress tolerance and plant growth.

### *In silico* analysis of *PpBAGs* gene expression

At the level of transcription, the plant *BAG* gene family has been associated with processes involving PCD, such as development and responses to biotic or abiotic stresses. In order to understand the expression pattern of *P. patens BAG* genes in different tissues, we performed an *in silico* analysis of the expression profile of *BAG* genes from *P. patens* using microarray data and RNA-seq based expression data from *P. patens*, available at Genevestigator (Hruz *et al*., 2008), *P. patens* eFP Browser (Ortiz-Ramírez *et al*., 2016) and PEATmoss (Perroud *et al*., 2018).

Under non-stress conditions, *PpBAG* genes were expressed during specific stages of the plant life cycle, but their expression pattern varied significantly between the different genes (Fig. 6a). Interestingly, we found that the expression of *PpBAGs* in different plant developmental stages was similar between genes encoding proteins that shared the same predicted subcellular localization and structural organization. *PpBAG1, PpBAG3* and *PpBAG8* were found to be specifically expressed in protoplasts and in the sporophytic and gametophytic phases of the moss life cycle. These three proteins belong to the same phylogenetic group and are predicted to localize in the nucleus. *PpBAG5* expression was slightly higher during the reproductive developmental phase compared to the vegetative developmental phase, and *PpBAG4* was more expressed in protonemata, being both PpBAG4 and PpBAG5 proteins with a UBL domain like their human BAG counterparts. *PpBAG6* was downregulated during reproductive development whereas *PpBAG7, PpBAG9* and *PpBAG10* did not show significant changes in their expression pattern during development.

**Fig. 6.**
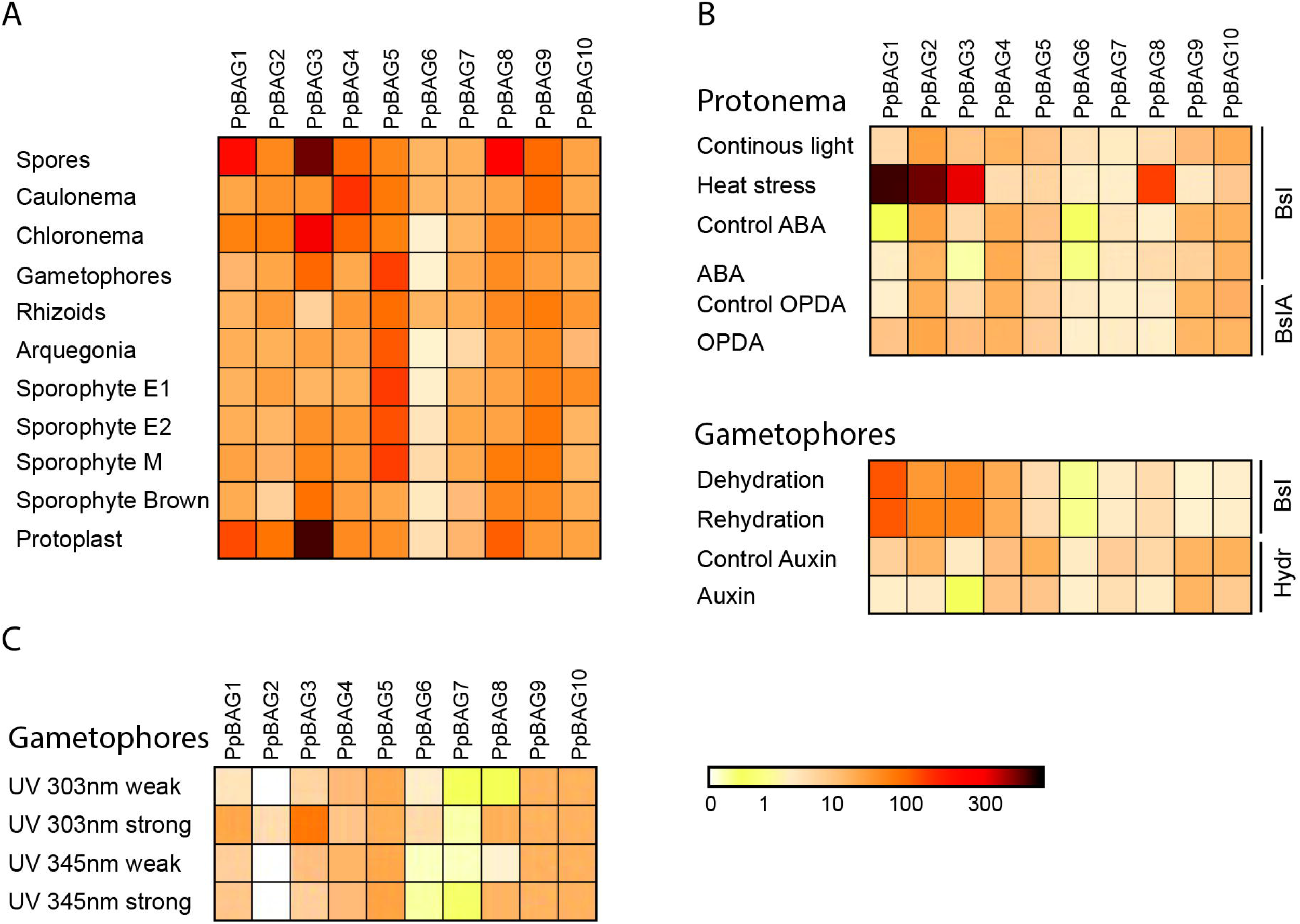
**a** Tissue-specific expression of *P. patens BAG* gene family from publicly available RNaseq data. Tissues and stages of development are as described in Ortiz-Ramirez et al. (2016). Sporophyte 1 (S1): sporophytes collected 5–6 days after fertilisation (AF), Sporophyte 2 (S2): Sporophytes collected 9–11 days AF, Sporophytes 3 (S3): Sporophyte collected 18–20 days AF and Sporophyte M (SM): Sporophytes collected 28–33 days AF. **b** Comparison of RNA-seq gene expression of *P. patens* BAG genes exposed to different hormonal treatments or stress conditions. Abbreviations: Bsl (BCD solid); BslA (BCDA (ammonium) solid); hydr (hydroponic); OPDA (12-oxophytodienoic acid). **c** Comparison of microarray gene expression of *P. patens BAG* genes exposed to UV-B (weak 303 nm and strong 345 nm).

Next, the transcriptional response of *P. patens BAG* genes was analyzed under various abiotic stresses and hormonal treatments. Comparative expression profiles indicated that only heat stress, dehydration and rehydration treatments resulted in significant upregulation of specific *PpBAGs* transcriptional levels (*PpBAG1-3* and *PpBAG8*) (Fig. 6b). These findings are consistent with the fact that *PpBAG1-3* and *PpBAG8* were the only members of the *PpBAG* gene family harboring HSF regulatory elements in their promoter regions (Fig. 5 and Table S2), and were also highly upregulated in protonemata during different time points of exposure to 37°C in comparison to 24°C (Elzanati., et al 2020) together with several PpHsp70s (Fig. S1), suggesting a role in responses to heat.

In addition, the same genes, with the exception of *PpBAG8*, were moderately induced in gametophores in response to dehydration and rehydration, while most of the other members of the gene family (*PpBAG5-9*) were not induced or down regulated under these conditions. The expression of *P. patens BAG* genes was also analyzed in response to hormonal treatments. *PpBAG* genes were unaffected by the ABA treatments (under the conditions studied), suggesting that *PpBAG1-3* and *PpBAG8* responded to heat and dehydration expression in an ABA-independent manner. However, *PpBAG1* and *PpBAG3* transcript levels were slightly increased by treatment with OPDA (50 μM OPDA and incubated for 6 h before harvesting), while *PpBAG3* was slightly repressed by ABA and auxin. Interestingly, a TGA regulatory element is present in the *PpBAG3* promoter region, suggesting a role for this gene in response to auxin.

Finally, microarray results under broad-band UV-B (303 nm) revealed *PpBAG1, PpBAG3* and *PpBAG8* up-regulation in gametophores (Fig. 6c), which was consistent with a previous gene expression analysis by UV-B radiation performed by Wolf *et al*. (2010). Contrastingly, in *A. thaliana* the expression of *AtBAG2, AtBAG3* (in shoots), and *AtBAG6* (in roots) was moderately down-regulated in response to UV-B (Nawkar *et al*., 2017).

### The protein-protein interaction network of *P. patens* BAGs

In order to gain insight into the *P. patens* BAG interactome, possible interacting partners were searched for each PpBAG protein using the STRING database, version 8 (Jensen *et al*., 2009). The predicted networks for the PpBAG proteins were mainly obtained based on text-mining and co-expression analysis (Fig. 7). Twenty proteins were found to potentially interact with five PpBAGs (PpBAG1, PpBAG3-PpBAG5 and 9), whereas no partners were predicted for the other five members of the PpBAG protein family. PpBAG4 and PpBAG5 were predicted to interact with a STUB1-like protein (CHIP, Pp3c14_2850V3.1), a chaperone-associated E3 ubiquitin ligase which can ubiquitinate chaperone-bound nonnative proteins and target them for degradation by the ubiquitin proteasome system (UPS) or selective autophagy under heat stress (Zhou *et al*., 2014). PpBAG1 and PpBAG3-5 may interact with specific HSP70 proteins (Pp1s91_109V6.1 and Pp1s29826V6.1). PpBAG4-5 also could also interact with SNF7 like protein, a key operator in the endosomal sorting complex required for transport (ESCRT). PpBAG1 and PpBAG3 are predicted to interact with other members of the same protein family (PpBAG4 and PpBAG5). Furthermore, PpBAG9 was also predicted to interact with PCD6 like proteins (Pp1s363_9V6.1 and Pp1s403_53V6.1), which mediate endosomal protein trafficking and turnover through the ESCRT/MVB pathway. In addition, PpBAG9 was predicted to interact with proteins with homology to SEC31 (Pp1s171_148V6.1, Pp1s281_108V6.1, and Pp1s171_149V6.1), a component of the coat protein complex I (COPII) which promotes the formation of transport vesicles from the endoplasmic reticulum (ER) (Ito *et al*., 2014). Overall, this data supports the involvement of some members of the *PpBAG* gene family in the response to heat stress and other responses involving the function of HSP70. Particularly interesting are the findings of several interacting proteins as members of the protein trafficking and turnover machinery, which suggest a role for BAG proteins in the regulation of the unfolded protein response.

**Fig. 7.**
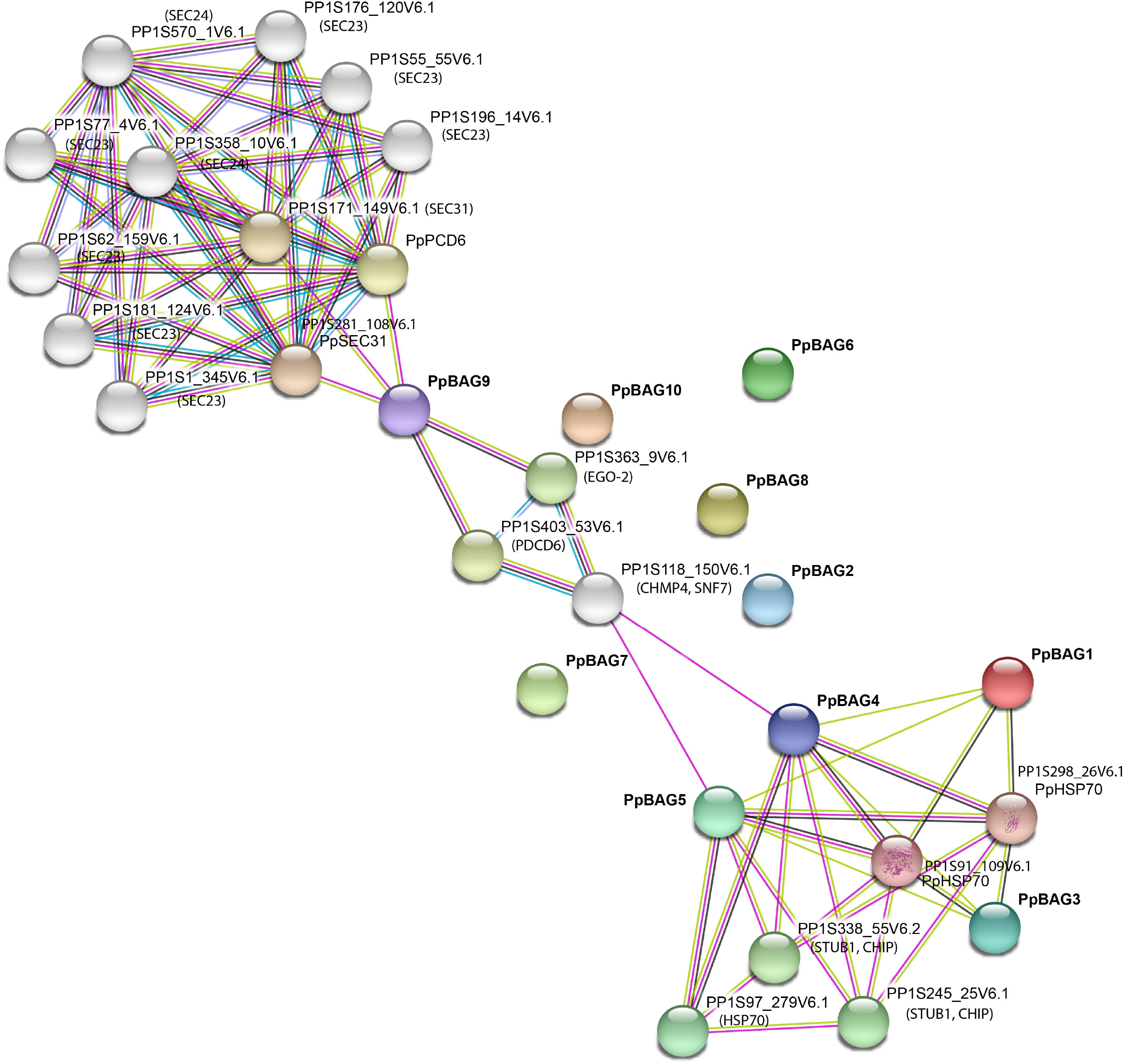
Protein-protein interaction network of *P. patens* BAG proteins. In the network generated by STRING version 11.0, each node represents a protein and each edge represents an interaction, colored by evidence type.

## Discussion

Organisms from bacteria to multicellular eukaryotes have the ability to induce PCD. In unicellular organisms, the cell is the organism *per se*, and it has been demonstrated that PCD benefits to kin instead of directly benefiting the organism itself (Vostinar *et al*., 2019). The fact that unicellular organisms are capable of undergoing PCD suggests that PCD has emerged early in life’s evolution and in multicellular organisms the evolution of this pathway has allowed cells to cooperate to increase the survival of the overall organism (Locato *et al*., 2018). In plants, PCD is a fundamental cellular process tightly controlled that eliminates specific cells under developmental or environmental stimuli (biotic and abiotic stresses). Despite the progress achieved on plant PCD, the molecular description of the key executioners and their control pathways remains poorly understood. In this sense, during the last decade investigations have revealed that BAG proteins are core regulators of plant PCD, participating in diverse pathways such as homeostasis, stress response and defense (Thanthrige *et al*., 2020). BAG proteins can act as nucleotide-exchange factors for Hsp70 family members that interact through the BD domain modulating the functions of these chaperones.

### Evolution of the *BAG* gene family in Viridiplantae

All plant BAGs contain the distinctive BAG domain located at the C terminus of the proteins, but different from the mammalian BAGs some plant BAGs contain a CaM-binding domain, named the IQ motif (Doukhanina *et al*., 2006), which appears to be plant specific and suggest a role in the Ca^2+^/CaM signaling pathways. In addition, the subcellular distribution of plant BAGs appears to be more diverse than in animals. While in animals these proteins are localized to the nucleus or the cytoplasm, in plants, they exhibit cytoplasmic localization or target different organelles such as mitochondria, vacuole, endoplasmic reticulum and nucleus (Thanthrige *et al*., 2020). Although experimental data is lacking, different subcellular localizations for PpBAGs have been predicted (Table 1). Taken together, these observations suggest that the plant BAGs may have evolved divergent roles compared with their animal counterparts.

Plant BAGs regulate cytoprotective processes from pathogen attack to abiotic stress and development. However, the mechanistic understanding of the different members of this family is rather limited and most of the knowledge on plant BAG proteins is restricted to angiosperms, mainly to the model eudicot *A. thaliana* (Cheng 2006; Doukhanina *et al*., 2006; Kang *et al*., 2005; Williams *et al*., 2010; Rana *et al*., 2012; Lee *et al*., 2016; Li *et al*., 2016; Wang *et al*., 2020). Extending these studies to other plant rich-species lineages in particular bryophytes (mosses, liverworts and hornworts), the sister clade of tracheophytes (vascular plants) as supported by recent analyses (Harris *et al*., 2020), may reveal novel paths/functions that have been undertaken during evolution (Rensing 2017).

Using different approaches including HHM based methods as previously used in *Arabidopsis* and rice (Doukhanina *et al*., 2006; Rana *et al*., 2012), we identified several *BAG* genes from different photosynthetic organisms including chlorophytes, charophytes and embryophytes. Like their mammalian counterparts, the common feature of all BAGs is the presence of the signature BD at the C-terminal region. Phylogenetic analysis performed in this study have included species which were not analyzed previously; chlorophytes (*Micromonas* sp. RCC299, *Micromonas pusilla* CCMP1545, *Coccomyxa subellipsoidea* C-169, *Dunaliella salina*), charophytes (*K. nitens*), bryophytes (*P. patens, M. polymorpha* and *S. fallax*), lycophytes (*S. moellendorffii*), amborella (*Amborella trichopoda*) and angiosperms (*Glycine max*) and categorized the *BAG* gene family into 2 subfamilies based on the presence of other conserved domains in addition to the BAG domain as previously described. Group I is represented by proteins with an UBL domain as observed in some mammalian BAG proteins like HsBAG1, is present along Viridiplantae; whereas Group II, characterized by the acquisition of the IQ motif in BAG proteins, is only present in embryophytes. Members of the chlorophyte species analyzed in this study contain single-copy *BAG* genes harbouring the UBL domain plus the BD domain, whereas *K. nitens* belonging to the charophytes, the ancestors of current terrestrial plants, has two *BAG* genes: one containing the UBL domain and one with a single BD. Gene expansion events were evident as plants evolved to more complex forms, ie. *G. max* with 24 *BAG* genes (Table S1).

The function of BAG proteins with a UBL domain (Group I) has been associated with the degradative function of Hsp70 chaperones. Mammalian BAG-1 and *S. pombe* Bag101 and Bag102 associate with the 26S proteasome via the N-terminal UBL domain and to Hsp70-type chaperones through the BD domain (Poulsen *et al*., 2016, Lüders *et al*., 2000), thus collaborating in the delivery of misfolded proteins to the proteasome leading to their degradation. In plants, AtBAG1 binds to endogenous Hsc70-4 through the BD domain, which specifically and cooperatively functions in the degradation of misfolded substrates and unimported chloroplast proteins in the cytosol in a proteasome-dependent manner (Lee *at al*., 2016; discussed below).

The fact that BAG proteins from Group II are restricted to embryophytes, suggests that the IQ signaling module evolved with the terrestrialization of plants. Although the IQ module is absent in BAG proteins from algae, it is present in other algal proteins. For example *Chara* myosin, a protein responsible for fast cytoplasmic streaming exhibits six IQ motifs for CaM binding in the motor domain (Morimatsu *et al*., 2000). In plants, the IQ motif is present in myosins, in CaM-binding transcriptional activators (CAMTAs), in cyclic nucleotide-gated channels (CNGCs), in addition to BAG proteins, but there are few reports regarding its functional role in these proteins. In AtCNGC20, a salt-regulated channel expressed in guard cells and in roots, the IQ motif binds CaM in a Ca^2+^-dependent manner at the plasma membrane which directs the C-terminus of CNGC20 into the nucleus (Fischer *et al*., 2013). However, the IQ motif exhibited by BAG proteins preferentially binds the calcium-free state of CaM (ApoCaM). The IQ motif of AtBAG5 binds to ApoCaM and the BAG domain binds to Hsc70, and in the presence of Ca^2+^ there is a competitive relationship between CaM and Hsc70 in binding to AtBAG5, which induce the release of Hsc70 by disrupting AtBAG5/Hsc70 binding (Li *et al*., 2016). AtBAG5 is localized to the mitochondria, and a model was proposed were elevated mitochondrial Ca2^+^ could induce the release of Hsp70 by disrupting AtBAG5/Hsp70 binding, leading to an inhibition of PCD progression and therefore attenuation of leaf senescence (Li *et al*., 2016). AtBAG6 also interacts with AtCaMs via the IQ motif in the absence of Ca^2+^, but both the CaM-binding IQ motif and the BAG domain are required for AtBAG6-mediated cell death in yeast and plants under oxidative stress (Kang *et al*., 2006). AtBAG7 also exhibits an IQ-motif but its functional role remains unknown.

The intron analysis performed in this study showed that genes belonging to Group IA present conserved intron-exon structures, whereas *BAG* genes of the Group IB and Group II exhibit different intron-exon structures and different numbers of exons. This suggests that groups of *BAG* genes derived from different ancestral genes, and the conserved intron phases observed in genes of Group IA (i.e., those located in the middle portion and in the of the C-terminal domain) indicate stability during evolution.

### *PpBAGs* are expressed during all stages of *P. patens* albeit a distinct set of genes respond to abiotic cues and OPDA treatment

Previous reports showed that plant BAG proteins act as co-chaperones regulating cell signaling, growth and development, as environmental stress responses (Doukhanina *et al*., 2006; Kim *et al*., 2011). Using an *in silico* approach we analyzed the transcriptional signature of *PpBAGs* during *P. patens* development and in response to different hormones and abiotic stresses. All *PpBAGs* were expressed during the different developmental stages of *P*. *patens* life cycle albeit *PpBAG1, PpBAG3* and *PpBAG8* exhibited the highest expression levels in spores and protoplasts. The expression signature in these tissues suggests a role in spore/protoplast germination and cell proliferation/elongation. *PpBAG3* showed the highest expression level among *PpBAGs* in chromonemata, spores, mature sporophyte and protoplasts, whereas *PpBAG5* exhibited the highest expression of all *PpBAGs* during sporophyte development (Fig. 6).

Plant hormones play important roles not only in growth and developmental processes but also in response to biotic and abiotic stresses. *In silico* analysis indicated that ABA induces expression of *PpBAG1* and *PpBAG8* at moderate levels. Search of *PpBAGs* expression profiles in different RNAseq data in *P. patens* treated with ABA at early time points (30, 60, and 180 min; 10 μM ABA or 1h, 100 μM) did not report changes in the expression level of these genes (Stevensson *et al*., 2016; Arif *et al*., 2019), but instead down-regulation of *PpBAG2, PpBAG3* and up-regulation of *PpBAG4* were detected in protonemata treated for 12h with 10 μM ABA (Shinozawa., et al 2019). Previous studies showed that the expression of Arabidopsis *BAGs* is regulated by ABA, which is important for adaptive responses to various environmental stresses. For example, *AtBGA6* is up-regulated by ABA, H202 and heat (Doukhanina et al. 2006; Kang et al. 2006). Furthermore, the expression of *PpBAG1-3* and *PpBAG8* was up-regulated in response to heat (Fig. 6 and Fig. S1) and *PpBAG1* and *PpBAG3* were also induced in response to OPDA treatment (*P. patens* produces OPDA but not JA). Interestingly, OPDA activates a COI1-independent pathway conserved in streptophyte plants that regulates plant thermotolerance genes which were likely crucial for plants successful land colonization (Monte *et al*., 2020). A recent study of the transcriptomic response of *P. patens* protonemata to heat (Elzanati *et al*., 2020) showed the up-regulation of genes involved in 12-oxo-phytodienoic acid (OPDA) synthesis, including lipoxygenase (12-LOX), allene oxide synthase (AOS) and allene oxide synthase (AOC). Treatment with these oxylipins protects *Marchantia* tissues against heat stress (Monte *et al*., 2020). It would be interesting to explore whether treatment with OPDA induces thermotolerance in *P. patens* tissues and analyze its relation with BAGs.

The presence of conserved stress-responsive cis-elements, such as DRE, ABRE, MYC, HSE, and MYB-binding sites, was confirmed in the promoter regions of *P. patens BAG* genes (Table S2, Fig. 5); suggesting that these elements are important for the regulation of their expression.

Several *PpBAGs* were downregulated by auxin namely, *PpBAG1-3, PpBAG5* and *PpBAG10*. Interestingly, *PpBAG1-3* were shown to be up-regulated under stress conditions, thus the pattern of downregulation in response to auxin, a major promoter of growth, suggests a mechanism of growth-defense tradeoff response.

### The potential role of PpBAGs under abiotic and biotic stresses: the link to autophagy?

Plant BAG proteins, like their mammalian counterparts, have been shown to regulate apoptosis-like processes, linking BAG proteins with plant PCD pathways under various abiotic and biotic stresses. *PpBAG1-3* and *PpBAG8* are induced by heat (Fig. 6), belong to the same phylogenetic group, and exhibit a similar modular structure harboring an IQ motif, in addition to the BD domain and a nuclear localization signature. In *Arabidopsis*, AtBAG7 (which harbors a NLS), is the unique *BAG* member constitutively localized in the endoplasmic reticulum and plays a central regulatory role in the induced Unfolded Protein Response (UPR) by heat (Williams *et al*., 2010). During thermal stress, AtBAG7 is proteolytically processed and translocated to the nucleus where it interacts with the transcription factor WRKY29 to induce the expression of genes coding for chaperones involved in stress tolerance (Li et al., 2017). Interestingly, a recent study of the transcriptional response of *P. patens* protonemata to heat (Elzanati *et al*., 2020) showed the up-regulation of *PpBAG1-3* and *PpBAG8*, together with several *HSP70s, WRKY, CHIP, SEC23, SEC24*, and several autophagic genes (*ATGs*) such as *ATG8s* and the selective autophagic marker *NBR1* (Fig. S1). The up-regulation of several members of the autophagic pathway in relation to *BAGs* needs to be further investigated. It has been shown that NBR1 and CHIP mediate two distinct but complementary anti-proteotoxic pathways, where NBR1-mediated autophagy targets ubiquitinated protein aggregates most likely derived from denatured or damaged nonnative proteins generated by heat, and CHIP mediating the degradation of nonnative proteins by the proteasome 26S (Zhou *et al;* 2014). It is unknown whether AtBAG7 is involved in the autophagic response to heat stress. However, the cleavage of AtBAG6 by an aspartyl protease (caspase-1 like) triggers autophagy and confers resistance to the necrotrophic fungal pathogen *B. cinerea* (Li *et al*., 2016). Noteworthy, *HsBAG3* expression is induced under acute oxidative or proteasomal stress and during cell aging and is inversely related to *HsBAG1* expression (known as the *BAG1-BAG3* Expression Shift). The Hs*BAG3* expression shift ensures sustained intracellular proteostasis by recruiting the selective macroautophagy pathway in association with HSP70, HSPB8 and the autophagy receptor p62 which specifically targets aggregation-prone proteins to autophagic degradation (Minoia et al., 2014). Furthermore, the presence of two IPV (Ile-Pro-Val) motifs in HsBAG3 are indispensable to regulate the interaction with the small heat-shock proteins (sHsp) sHsp6 and sHsp8 (Carra *et al*., 2008; Fuchs *et al*., 2009). Sequences analysis revealed the presence of one IPV motif in PpBAG1-3 and two motifs in PpBAG8 (Fig. 2).

BAG proteins have also been associated with responses to pathogens. As mentioned, AtBAG6 is involved in the autophagy pathway, particularly as part of the plant defense response in the infection to necrotrophic pathogens (Li *et al*., 2016). AtBAG6 is cleaved by caspase-like proteases and triggers autophagy in the host to limit fungal colonization and confer basal resistance to necrotrophic fungus *Botrytis cinerea* (Li *et al*., 2016). This fungal pathogen causes severe necrosis in *P. patens* leading to plant maceration and cell death with hallmarks of PCD (Ponce de León *et al*., 2012). However, no induction of *PpBAGs* genes in response to *Botrytis* was observed in the available microarray and RNA-seq data. Given that PpBAG1, PpBAG3 and PpBAG9 deduced proteins have a predicted caspase-1 cleavage site downstream of the BAG domain, and that these proteins clustered with AtBAG6 in the phylogenetic analysis (Fig. 2 and 1, respectively), it is likely that these particular members of the *P. patens* BAG family could play a role in pathogen defense. Further studies are required to determine whether these proteins can be proteolytically activated to induce autophagic cell death and plant resistance responses.

### PpBAG4 and PpBAG5 as candidates in protein quality control via the proteasome system

Our analysis revealed the existence of ten genes encoding BAG proteins in the *P. patens* genome, which were categorized into two subfamilies with PpBAG4, PpBAG5 and PpBAG10 belonging to Group I and PpBAG1-3 and PpBAG6-9 to Group II. PpBAG4 and PpBAG5 contain a conserved UBL domain (Fig. 3), as also observed in AtBAG1-4 and in the mammalian HsBAG-1 protein (Fig. 2), and differ in their expression pattern with PpBAG4 mainly expressed in protonemata and PpBAG5 during the reproductive stages of *P. patens* life cycle (Fig. 5). A functional role as a co-chaperone of Hsc70-4-mediated proteasomal degradation of misfolded and unimported plastid proteins in the cytosol has been assigned to AtBAG1, executed through the interaction with Hsc70-4 through the BD domain (Lee *et al*., 2016), and putatively with CHIP (although this possibility needs to be experimentally addressed) through the UBL domain, as known for HsBAG-1 (Demand *et al*., 2001). Comparative sequence alignment between mammals and plants showed that most residues required for Hsp70 binding in mammals are conserved in *P. patens* BAG proteins (Fig. 2B). Therefore, it is reasonable to speculate that PpBAGs bind Hsp70 in a manner similar to that of their mammalian counterparts. Interestingly, the interactome analysis predicted interactions of both PpBAG4 and PpBAG5 with both Hsp70s and a CHIP protein (Fig. 7), highlighting their possible role as Hsp70 co-chaperones involved in protein quality control, providing a link to the proteasome system. In agreement with this putative function, a cytoplasmic localization has been predicted for PpBAG4 and PpBAG5 proteins, in similar fashion as shown for AtBAG1–3 which are mainly localized in the cytosol, and AtBAG4 localized in both the cytosol and the nucleus (Lee *et al*., 2016), where the ubiquitin-proteasome system is localized.

### PpBAG as candidate for mitochondrial Ca^2+^ uptake

PpBAG9 is predicted to localize in the mitochondria. Besides the BD domain and IQ motif, it also has an EF-Hand motif near the C-terminal region, a novel feature that has not been described so far for members of this family. Mitochondrial EF-hand proteins are central to the proper decoding Ca^2+^ fluctuations to control both metabolism and stress responses (Hajnóczky *et al*., 2014). Interestingly, the mammalian apoptosis-linked gene 2 (ALG-2/PCDC-6), is a protein with a penta EF domain. Ca^2+^ binding to its PEF domain results in conformational changes that increase the hydrophobicity of the protein and consequently the interaction with its protein targets, which have been reported to indirectly promote the cell death pathway at multiple steps (Maki *et al*, 2019). More research is needed to underpin the role of the EF domain in PpBAG9 which appears to be a specie-specific feature.

In summary, we presented the first comprehensive analysis of the *BAG* gene family in the bryophyte *Physcomitrium patens*, a non flowering plant. Phylogenetic analysis revealed the presence of two distinct Groups of plant *BAG* genes, which differ from each other in structural features, exon/intron organization, and phase pattern. Whereas Group I harbours the more ancestral BAGs, Group II is only present in land plants and reported phenotypes and expression patterns for proteins of Group II suggest their role in plant adaptation to land (dehydration habitats, variations in temperature, ultraviolet radiation, high light intensities and pathogen infection). Based on our study, candidate genes from this group (*PpBAG1-3* and *PpBAG8*) were identified in *P. patens* for further functional characterization under abiotic stresses especially under heat stress and their possible role in connection to autophagy. Additionally, caspase-1 predicted sites PpBAG1, PpBAG3-5 and PpBAG9 invite to explore the processing of these genes for cell death/cytoprotective activity. In addition, we identified PpBAG9 which harbors a *P. patens* BAG specific feature, an EF-domain not reported in BAG proteins previously. Our study reinforces the notion that expanding in the use of other model organisms such as Bryophytes will provide progress in our understanding of fundamental aspects of plant responses to stress.

## Abbreviations

BD: BAG Domain
PCD: Programmed Cell Death
BAG: Bcl-2-Associated athanogene
UBL: Ubiquitin-like
CaM: Calmodulin
UPS: Ubiquitin Proteasome System

## Acknowledgements

This work was supported by grants from Agencia Nacional de Investigación e Innovación (ANII) FCE_2018_148590, Agencia Nacional de Promoción Científica y Tecnológica, Argentina (FONCYT-PICT-2016-0497) and Consejo Nacional de Investigaciones Científicas y Técnicas (CONICET).

## Author contributions

A.C and L.S. designed the study. L.S., C.R and A.C analyzed the data. A.C and L.S. wrote the manuscript with valuable input from R.L and S.V. All authors have read and approved this manuscript.

---

**Table S1** List of *BAG* genes used in this study.

**Table S2** PpBAGs promoter cis-acting regulatory elements analyzed with PlantCARE.

**Fig. S1** Up-regulated *BAG*, Autophagy (*ATG*), *CHIP, SEC, HSP70s, sHSPs and HSP90s* genes in *P. patens* protonemata under high heat stress (37°C) compared to control conditions (24°C). 1h, 6h, 12h and 24h stand for the time grown at the respective temperature. The expression data was selected from the RNA-Seq study performed by Elzanati, et al (2020). Genes IDs: *PpATG1a (Pp3c19_5230), PpATG11a (Pp3c19_12090), PpATG13b (Pp3c2_10630), PpVPS15b (Pp3c9_3860), PpATG9b (Pp3c2_1820), PpATG4b (Pp3c7_280), PpATG8e (Pp3c14_15040), PpATG8f (Pp3c14_15141), PpNBR1a (Pp3c7_8990), PpNBR1b (Pp3c11_16970)*.

## References

Arif, M. A., Hiss, M., Tomek, M., Busch, H., Meyberg, R., Tintelnot, S., Reski, R., Rensing, S. A., & Frank, W. (2019). ABA-Induced Vegetative Diaspore Formation in *Physcomitrella patens*. Frontiers in plant science, 10, 315. https://doi.org/10.3389/fpls.2019.00315

Ashton NW, Grimsley NH, Cove DJ (1979). Analysis of gametophytic development in the moss, *Physcomitrella patens,* using auxin and cytokinin resistant mutants. Planta. 144(5):427–435. doi:10.1007/BF00380118

Balakireva, A. V., & Zamyatnin, A. A., Jr (2019). Cutting Out the Gaps Between Proteases and Programmed Cell Death. Frontiers in plant science, 10, 704. https://doi.org/10.3389/fpls.2019.00704

Behl C. (2016). Breaking BAG: The Co-Chaperone BAG3 in Health and Disease. Trends in pharmacological sciences, 37(8), 672–688. https://doi.org/10.1016/j.tips.2016.04.007

Carra S, Seguin SJ, Lambert H, Landry J (2008) HspB8 chaperone activity toward poly(Q)-containing proteins depends on its association with Bag3, a stimulator of macroautophagy. J Biol Chem. 283(3):1437–44x.

Daneva, A., Gao, Z., Van Durme, M., & Nowack, M. K. (2016). Functions and Regulation of Programmed Cell Death in Plant Development. Annual review of cell and developmental biology, 32, 441–468. https://doi.org/10.1146/annurev-cellbio-111315-124915

DeLano, W.L. The PyMOL User’s Manual (2002) DeLano Scientific, San Carlos, CA, USA.

Demand J, Alberti S, Patterson C, Höhfeld J. (2001) Cooperation of a ubiquitin domain protein and an E3 ubiquitin ligase during chaperone/proteasome coupling. Curr Biol. 2001 Oct 16;11(20):1569–77. doi: 10.1016/s0960-9822(01)00487-0

Doolittle WF. (1978). Genes in piece: Were they ever together? Nature (London) 272:581–582.

Doukhanina EV, Chen S, van der Zalm E, Godzik A., Reed J, Dickman MB. (2006). Identification and functional characterization of the BAG protein family in *Arabidopsis thaliana*. J. Biol. Chem. 281, 18793–18801.

Elzanati, O., Mouzeyar, S., & Roche, J. (2020). Dynamics of the Transcriptome Response to Heat in the Moss, Physcomitrella patens. International journal of molecular sciences, 21(4), 1512. https://doi.org/10.3390/ijms21041512

Emanuelsson O, Nielsen H, Brunak S, von Heijne G. (2000) Predicting subcellular localization of proteins based on their N-terminal amino acid sequence. J Mol Biol. 300(4):1005–1016. doi:10.1006/jmbi.2000.3903

Fernandez-Pozo N, Haas FB, Meyberg R, et al. (2020) PEATmoss *(Physcomitrella* Expression Atlas Tool): a unified gene expression atlas for the model plant *Physcomitrella patens*. Plant J. 102(1):165–177. doi:10.1111/tpj.14607

Fischer, C., Kugler, A., Hoth, S., & Dietrich, P. (2013). An IQ domain mediates the interaction with calmodulin in a plant cyclic nucleotide-gated channel. Plant & cell physiology, 54(4), 573–584. https://doi.org/10.1093/pcp/pct021

Froesch, B. A., Takayama, S. and Reed, J. C. (1998). BAG-1L protein enhances androgen receptor function. J. Biol. Chem. 273, 11660–11666.

Fuss, J., Liegmann, O., Krause, K., and Rensing, S. A. (2013). Green targeting predictor and ambiguous targeting predictor 2: the pitfalls of plant protein targeting prediction and of transient protein expression in heterologous systems. New Phytol. 200:1022.

Gilbert W (1987) Cold Spring Harb Symp Quant Biol. 52:901–5.

Gilbert W, de Souza SJ, Long M (1997). Proc Natl Acad Sci U S A. Jul 22; 94(15):7698–703.

Hajnóczky, G., Booth, D., Csordás, G., Debattisti, V., Golenár, T., Naghdi, S., Niknejad, N., Paillard, M., Seifert, E. L., & Weaver, D. (2014). Reliance of ER-mitochondrial calcium signaling on mitochondrial EF-hand Ca^2+^ binding proteins: Miros, MICUs, LETM1 and solute carriers. Current opinion in cell biology, 29, 133–141. https://doi.org/10.1016/j.ceb.2014.06.002

Harris BJ, Harrison CJ, Hetherington AM, Williams TA. (2020) Phylogenomic Evidence for the Monophyly of Bryophytes and the Reductive Evolution of Stomata. Curr Biol. Jun 8;30(11):2001–2012.e2. doi: 10.1016/j.cub.2020.03.048

Hruz, T., Laule, O., Szabo, G., Wessendorp, F., Bleuler, S., Oertle, L., et al. (2008). Genevestigator v3: a reference expression database for the meta-analysis of transcriptomes. Adv. Bioinformatics 2008:420747.

Hu B, Jin, Guo AJ, Zhang H, Luo J, Gao G. (2015). GSDS 2.0: an upgraded gene feature visualization server. Bioinformatics, 31(8):1296–1297.

Ito Y, Uemura T, Nakano A. (2014) Formation and maintenance of the Golgi apparatus in plant cells. Int Rev Cell Mol Biol. 310:221–87. doi: 10.1016/B978-0-12-800180-6.00006-2

Kabbage M., Dickman M. B. (2008). The BAG proteins: a ubiquitous family of chaperone regulators. Cell Mol. Life Sci. 65 1390–1402.

Kang CH, Jung WY, Kang YH, et al. (2006) AtBAG6, a novel calmodulin-binding protein, induces programmed cell death in yeast and plants. Cell Death Differ. 13, 84–95.

Kenrick, P. (2017). How land plant life cycles first evolved. Science 358, 1538–1539.

Kim MC, Panstruga R, Elliott C, et al. (2002) Calmodulin interacts with MLO protein to regulate defence against mildew in barley. Nature. 416(6879):447–451.

Li, L., Xing, Y., Chang, D., Fang, S., Cui, B., Li, Q., Wang, X., Guo, S., Yang, X., Men, S., & Shen, Y. (2016). CaM/BAG5/Hsc70 signaling complex dynamically regulates leaf senescence. Scientific reports, 6, 31889. https://doi.org/10.1038/srep31889

Li Y, Kabbage M, Liu W, Dickman MB. (2016). Aspartyl Protease-Mediated Cleavage of BAG6 Is Necessary for Autophagy and Fungal Resistance in Plants. Plant Cell. 28(1):233–247. doi:10.1105/tpc.15.00626

Li Y, Williams B, Dickman M. (2017) *Arabidopsis* B-cell lymphoma2 (Bcl-2)-associated athanogene 7 (BAG7)-mediated heat tolerance requires translocation, sumoylation and binding to WRKY29. New Phytol. 214(2):695–705. doi:10.1111/nph.14388

Lee, D. W., Kim, S. J., Oh, Y. J., Choi, B., Lee, J., & Hwang, I. (2016). *Arabidopsis* BAG1 Functions as a Cofactor in Hsc70-Mediated Proteasomal Degradation of Unimported Plastid Proteins. Molecular plant, 9(10), 1428–1431. https://doi.org/10.1016/j.molp.2016.06.005

Lüders, J., Demand, J., & Höhfeld, J. (2000). The ubiquitin-related BAG-1 provides a link between the molecular chaperones Hsc70/Hsp70 and the proteasome. The Journal of biological chemistry, 275(7), 4613–4617. https://doi.org/10.1074/jbc.275.7.4613

Maki M, Kitaura Y, Satoh H, Ohkouchi S, Shibata H. (2002) Structures, functions and molecular evolution of the penta-EF-hand Ca2+-binding proteins. Biochim Biophys Acta. 1600(1-2):51–60. doi: 10.1016/s1570-9639(02)00444-2. PMID: 12445459

Minoia M, Boncoraglio A, Vinet J, Morelli FF, Brunsting JF, Poletti A, Krom S, Reits E, Kampinga HH, Carra S. (2014) BAG3 induces the sequestration of proteasomal clients into cytoplasmic puncta: implications for a proteasome-to-autophagy switch. Autophagy. Sep;10(9):1603–21. doi: 10.4161/auto.29409

Monte, I., Kneeshaw, S., Franco-Zorrilla, J. M., Chini, A., Zamarreño, A. M., García-Mina, J. M., & Solano, R. (2020). An Ancient COI1-Independent Function for Reactive Electrophilic Oxylipins in Thermotolerance. Current biology: CB, 30(6), 962–971.e3. https://doi.org/10.1016/j.cub.2020.01.023

Morimatsu, M., Nakamura, A., Sumiyoshi, H., Sakaba, N., Taniguchi, H., Kohama, K., & Higashi-Fujime, S. (2000). The molecular structure of the fastest myosin from green algae, Chara. Biochemical and biophysical research communications, 270(1), 147–152. https://doi.org/10.1006/bbrc.2000.2391

Nawkar, G. M., Maibam, P., Park, J. H., Woo, S. G., Kim, C. Y., Lee, S. Y., & Kang, C. H. (2017). In silico study on *Arabidopsis* BAG gene expression in response to environmental stresses. Protoplasma, 254(1), 409–421. https://doi.org/10.1007/s00709-016-0961-3

Nguyen HD, Yoshihama M, Kenmochi N. (2006) Phase distribution of spliceosomal introns: implications for intron origin. BMC Evol Biol. 6:69.

Ortiz-Ramírez, C., Hernandez-Coronado, M., Thamm, A., Catarino, B., Wang, M., Dolan, L., et al. (2016). A transcriptome atlas of *Physcomitrella patens* provides insights into the evolution and development of land plants. Mol. Plant. 9, 205–220.

Perroud PF, Haas FB, Hiss M, et al. (2019) The *Physcomitrella patens* gene atlas project: large-scale RNA-seq based expression data [published correction appears in Plant J. 95(1):168–182.

Ponce De León I, Schmelz EA, Gaggero C, Castro A, Álvarez A, Montesano M. (2012) *Physcomitrella patens* activates reinforcement of the cell wall, programmed cell death and accumulation of evolutionary conserved defence signals, such as salicylic acid and 12-oxo-phytodienoic acid, but not jasmonic acid, upon *Botrytis cinerea* infection. Mol Plant Pathol. 13(8):960–974. doi:10.1111/j.1364-3703.2012.00806.x

Poulsen, E. G., Kampmeyer, C., Kriegenburg, F., Johansen, J. V., Hofmann, K., Holmberg, C., & Hartmann-Petersen, R. (2017). UBL/BAG-domain co-chaperones cause cellular stress upon overexpression through constitutive activation of Hsf1. Cell stress & chaperones, 22(1), 143–154. https://doi.org/10.1007/s12192-016-0751-z

Putkey, J. A., Kleerekoper, Q., Gaertner, T. R. & Waxham, M. N. (2003). J. Biol. Chem. 278, 49667–49670.

Rana, Rashid & Dong, Shinan & Ali, Zulfiqar & Khan, Azeem & Zhang, Hong. (2012). Identification and characterization of the Bcl-2-associated athanogene (BAG) protein family in rice. African J of Biotechnology. 11. 88–98.

Rensing, S. A., Lang, D., Zimmer, A. D., Terry, S., Salamov, S., Shapiro, H., et al. (2008). The Physcomitrella genome reveals evolutionary insights into the conquest of land by plants. Science 319, 64–69.

Rensing, S. A. (2018). Great moments in evolution: the conquest of land by plants. Curr. Opin. Plant Biol. 42, 49–54.

Rogozin IB, Carmel L, Csuros M, Koonin EV. (2012) Origin and evolution of spliceosomal introns. Biol Direct. 7:11.

Shinozawa, A., Otake, R., Takezawa, D., Umezawa, T., Komatsu, K., Tanaka, K., Amagai, A., Ishikawa, S., Hara, Y., Kamisugi, Y., Cuming, A. C., Hori, K., Ohta, H., Takahashi, F., Shinozaki, K., Hayashi, T., Taji, T., & Sakata, Y. (2019). SnRK2 protein kinases represent an ancient system in plants for adaptation to a terrestrial environment. Communications biology, 2, 30. https://doi.org/10.1038/s42003-019-0281-1

Sondermann H, Scheufler C, Schneider C, Hohfeld J, Hartl FU, Moarefi I. (2001) Structure of a Bag/Hsc70 complex: convergent functional evolution of Hsp70 nucleotide exchange factors. Science. 291(5508):1553–1557.

Stevenson, S. R., Kamisugi, Y., Trinh, C. H., Schmutz, J., Jenkins, J. W., Grimwood, J., Muchero, W., Tuskan, G. A., Rensing, S. A., Lang, D., Reski, R., Melkonian, M., Rothfels, C. J., Li, F. W., Larsson, A., Wong, G. K., Edwards, T. A., & Cuming, A. C. (2016). Genetic Analysis of *Physcomitrella patens* Identifies ABSCISIC ACID NON-RESPONSIVE, a Regulator of ABA Responses Unique to Basal Land Plants and Required for Desiccation Tolerance. The Plant cell, 28(6), 1310–1327. https://doi.org/10.1105/tpc.16.00091

Szklarczyk D, Gable AL, Lyon D, Junge A, Wyder S, Huerta-Cepas J, Simonovic M, Doncheva NT, Morris JH, Bork P, Jensen LJ, von Mering C. (2019) STRING v11: protein-protein association networks with increased coverage, supporting functional discovery in genome-wide experimental datasets. Nucleic Acids Res. Jan; 47:D607–613.

Tamura, K., Peterson, D., Peterson, N., Stecher, G., Nei, M., and Kumar, S. (2011). MEGA5: molecular evolutionary genetics analysis using maximum likelihood, evolutionary distance, and maximum parsimony methods. Mol. Biol. Evol. 28, 2731–2739.

Terada K, Mori M. (2000) Human DnaJ homologs dj2 and dj3, and bag-1 are positive cochaperones of hsc70. J Biol Chem. 275(32):24728–24734.

Thanthrige N, Jain S, Bhowmik SD, et al. (2020) Centrality of BAGs in Plant PCD, Stress Responses, and Host Defense. Trends Plant Sci. (20) 30127–8.

Wang, J., Yeckel, G., Kandoth, P. K., Wasala, L., Hussey, R. S., Davis, E. L., Baum, T. J., & Mitchum, M. G. (2020). Targeted suppression of soybean BAG6-induced cell death in yeast by soybean cyst nematode effectors. Molecular plant pathology, 21(9), 1227–1239. https://doi.org/10.1111/mpp.12970

Wilkins MR, Gasteiger E, Bairoch A, et al. (1999) Protein identification and analysis tools in the ExPASy server. Methods Mol Biol. 112:531–552. doi:10.1385/1-59259-584-7:531

Williams B, Kabbage M, Britt R, Dickman MB. (2010) AtBAG7, an *Arabidopsis* Bcl-2-associated athanogene, resides in the endoplasmic reticulum and is involved in the unfolded protein response. Proc Natl Acad Sci U S A. 107(13):6088–6093. doi:10.1073/pnas.0912670107

Reape TJ, Molony EM, McCabe PF. (2008) Programmed cell death in plants: distinguishing between different modes. J Exp Bot. 59(3):435–444. doi:10.1093/jxb/erm258

Vostinar AE, Goldsby HJ, Ofria C (2019) Suicidal selection: Programmed cell death can evolve in unicellular organisms due solely to kin selection. Ecol Evol. Aug; 9(16): 9129–9136.

Zhou J, Zhang Y, Qi J, et al. (2014) E3 ubiquitin ligase CHIP and NBR1-mediated selective autophagy protect additively against proteotoxicity in plant stress responses [published correction appears in PLoS Genet. 2014 Jun;10(6):e1004478]. PLoS Genet. 10(1):e1004116. doi:10.1371/journal.pgen.1004116

