## Supplementary figures and images for "Genome-wide identification, characterization and expression analysis of the Bcl-2 associated athanogene (BAG) gene family in *Physcomitrium patens*"

### Supplemental Figure 1

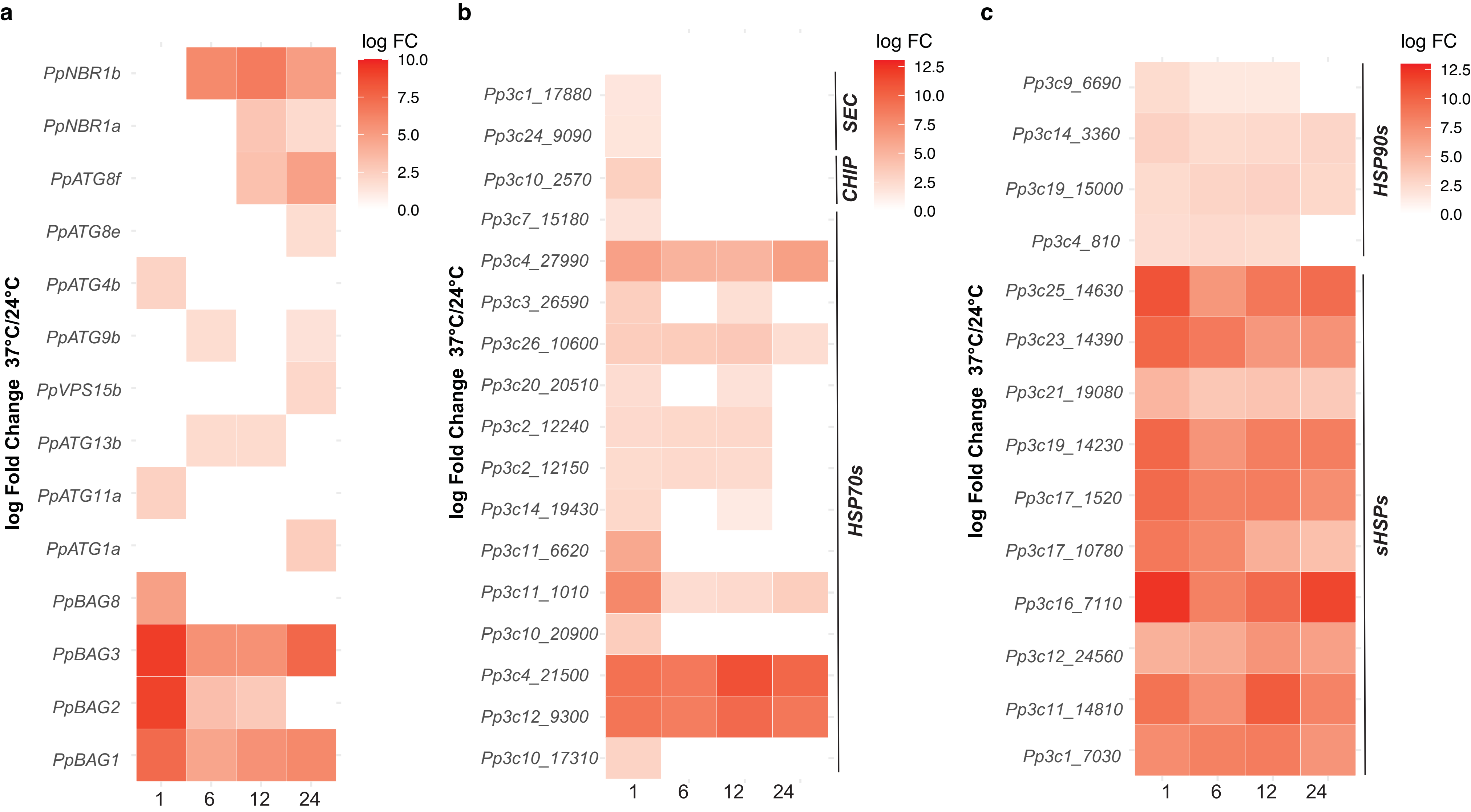
